# METAbolomics data Balancing with Over-sampling Algorithms (META-BOA): an online resource for addressing class imbalance

**DOI:** 10.1101/2022.04.21.489108

**Authors:** Emily Hashimoto-Roth, Anuradha Surendra, Mathieu Lavallée-Adam, Steffany A. L. Bennett, Miroslava Čuperlović-Culf

## Abstract

**Motivation:** Class imbalance, or unequal sample sizes between classes, is an increasing concern in machine learning for metabolomic and lipidomic data mining, which can result in overfitting for the over-represented class. Numerous methods have been developed for handling class imbalance, but they are not readily accessible to users with limited computational experience. Moreover, there is no resource that enables users to easily evaluate the effect of different over-sampling algorithms.

**Results:** METAbolomics data Balancing with Over-sampling Algorithms (META-BOA) is a web-based application that enables users to select between four different methods for class balancing, followed by data visualization and classification of the sample to observe the augmentation effects. META-BOA outputs a newly balanced dataset, generating additional samples in the minority class, according to the user’s choice of Synthetic Minority Over-sampling Technique (SMOTE), Borderline-SMOTE (BSMOTE), Adaptive Synthetic (ADASYN), or Random Over-Sampling Examples (ROSE). META-BOA further displays both principal component analysis (PCA) and t-distributed stochastic neighbor embedding (t-SNE) visualization of data pre- and post-over-sampling. Random forest classification is utilized to compare sample classification in both the original and balanced datasets, enabling users to select the most appropriate method for their analyses.

**Availability and implementation:** META-BOA is available at https://complimet.ca/meta-boa.

**Supplementary Information:** Supplementary material is available at *Bioinformatics* online.

## 1 INTRODUCTION

Metabolomics and lipidomics are emerging fields with unprecedented potential to elucidate novel pathogenic metabolic processes, producing diagnostic and treatment-relevant knowledge. Both “omics” disciplines generate high dimensional data. Yet classes of samples are often not equally balanced within datasets, presenting a notable challenge for machine learning strategies. While some machine learning models, such as support vector machines, are relatively robust to imbalanced datasets, most models are subject to overfitting, favouring the over-represented class. These challenges can bias overall analyses, contrary to the intent of an unbiased machine learning approach. Class imbalance can be addressed by under-sampling the majority class or over-sampling the minority class using a variety of computational algorithms (reviewed in (Kovács, 2019; Sharma, et al., 2022)). However, to our knowledge, there is no tool that enables researchers to rapidly perform and compare different over-sampling methodologies. METAbolomics data Balancing with Over-sampling Algorithms (META-BOA) is a user-friendly web-based application that generates balanced datasets, for both binary and multiclass datasets. Users can additionally evaluate each methodology through visual representations of unsupervised and supervised analyses, comprising of PCA and t-SNE data visualizations and random forest classification.

## 2 Implementation

META-BOA is an application, accessible through any web browser that, through over-sampling, generates balanced sample groups for both binary and multi-class datasets. META-BOA provides users with the option to select between four over-sampling methods: SMOTE, BSMOTE, ADASYN, and ROSE. These algorithms were chosen to maximize user alternatives: SMOTE randomly generates new synthetic samples within the known minority class samples without replication (Chawla, et al., 2002). BSMOTE generates synthetic samples on the borderline between majority and minority instances (Han, et al., 2005). ADASYN creates more samples in the neighborhood of minority samples that are in the vicinity of a larger number of the majority class cases. (He, et al., 2008). Finally, ROSE is a bootstrap-based approach that creates synthetic samples in the neighbourhood of minority class features (Lunardon, et al., 2014).

The META-BOA workflow is presented in Figure 1. The platform was coded in Python 3.7 and deployed with an R Shiny graphical user interface. To run META-BOA, the user selects a single .CSV file containing their imbalanced dataset. The user then specifies the following parameters:

1. Choice of over-sampling algorithm (default = SMOTE)
2. Normalization applied to data (default = no normalization)
3. Dimensionality reduction and visualization by PCA and t-SNE analysis (default = yes)

**Figure 1.**
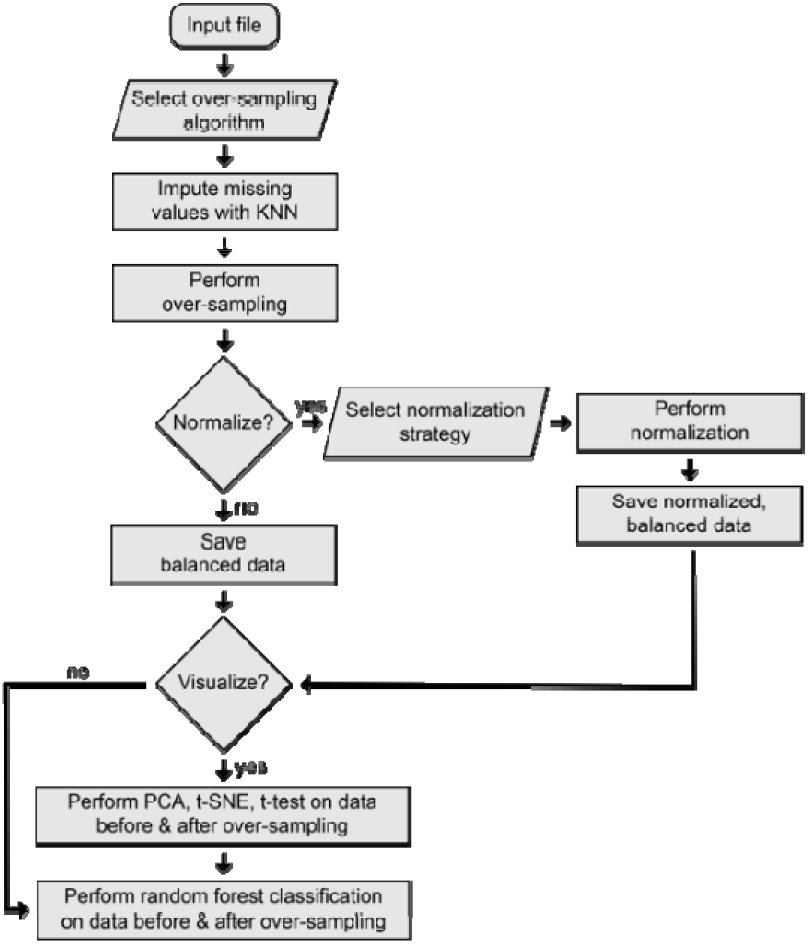
Outline of META-BOA workflow. Output is class balanced dataset with or without normalization.

Users can choose from four over-sampling algorithms: (1) SMOTE, (2) BSMOTE, (3) ADASYN, and (4) ROSE. Each option has been implemented using the *imblearn* package (version 0.8.0) (Lemaitre, et al., 2017). Data visualization and random forest classification provide a means to evaluate the effect of the selected over-sampling algorithm. META-BOA performs both PCA and t-SNE dimensionality reduction, evaluating linear and non-linear reduction, respectively. The random forest classification model is trained and tested with an 80/20 random split, and with 250 trees. These evaluations are intended to provide a preliminary analysis of the newly balanced dataset, while showing the effect of over-sampling on the visualization and classification results. Dimensionality reduction and classification are implemented using the Scikit-learn package (version 0.24.0) (Pedregosa, et al., 2011). Result files are made available for download immediately upon analysis completion, as a zipped file which includes a .CSV datafile and .SVG image files.

## 3 Conclusion

META-BOA is an open-access web-based application for balancing sample classes in metabolomics and lipidomics datasets, or any other type of data, using over-sampling algorithms. This tool will help facilitate future machine learning-based analyses and modeling of me-tabolomic and lipidomic data. We encourage users to implement METABOA in their data pre-processing pipelines, as well as test the different over-sampling algorithms available to evaluate their effects on their given datasets.

## Funding

This work was supported in part by RGPIN-2019-06796 to SALB from the Natural Sciences and Engineering Research Council of Canada (NSERC), as well as an NSERC CREATE Matrix Metabolomics Training grant to SALB and MLA, an operating grants AI-4D-102-3 to SALB and MCC from the National Research Council AI for Design Challenge Program, and an NSERC Discovery Grant to MLA. This research was enabled by support provided by Compute Ontario and Compute Canada with Resource Allocation to M.L.A. EHR received a NSERC CREATE Matrix Metabolomics Scholarship.

## Conflict of Interest

none declared.

## Notes

### Competing Interest Statement

The authors have declared no competing interest.

### Summary of Updates

hyphens were in wrong places in title and abstract.

