## Supplementary Material for "METAbolomics data Balancing with Over-sampling Algorithms (META-BOA): an online resource for addressing class imbalance"

| Figure S1 | Comparison of ROC curve results using four different over-sampling methods and the original dataset all with z-score normalization of log transformed data. |
| --- | --- |
| Figure S2 | Comparison of PCA results using four different over-sampling methods and the original dataset all with z-score normalization of log transformed data. |
| Figure S3 | Comparison of t-SNE results using four different over-sampling methods and the original dataset all with z-score normalization of log transformed data. |

Example datasets and detailed instructions for META-BOA are provided at: <https://complimet.ca/meta-boa/>.

**Figure S1.** Comparison of ROC curve results using four different over-sampling methods and original dataset all following z-score normalization of log transformed data. Comparison of Area Under the Curve (AUC) values for three samples shows that for this sample set SMOTE method leads to the highest classification accuracy. At the same time all over-sampling methods improved classification when compared with the imbalanced, original dataset.

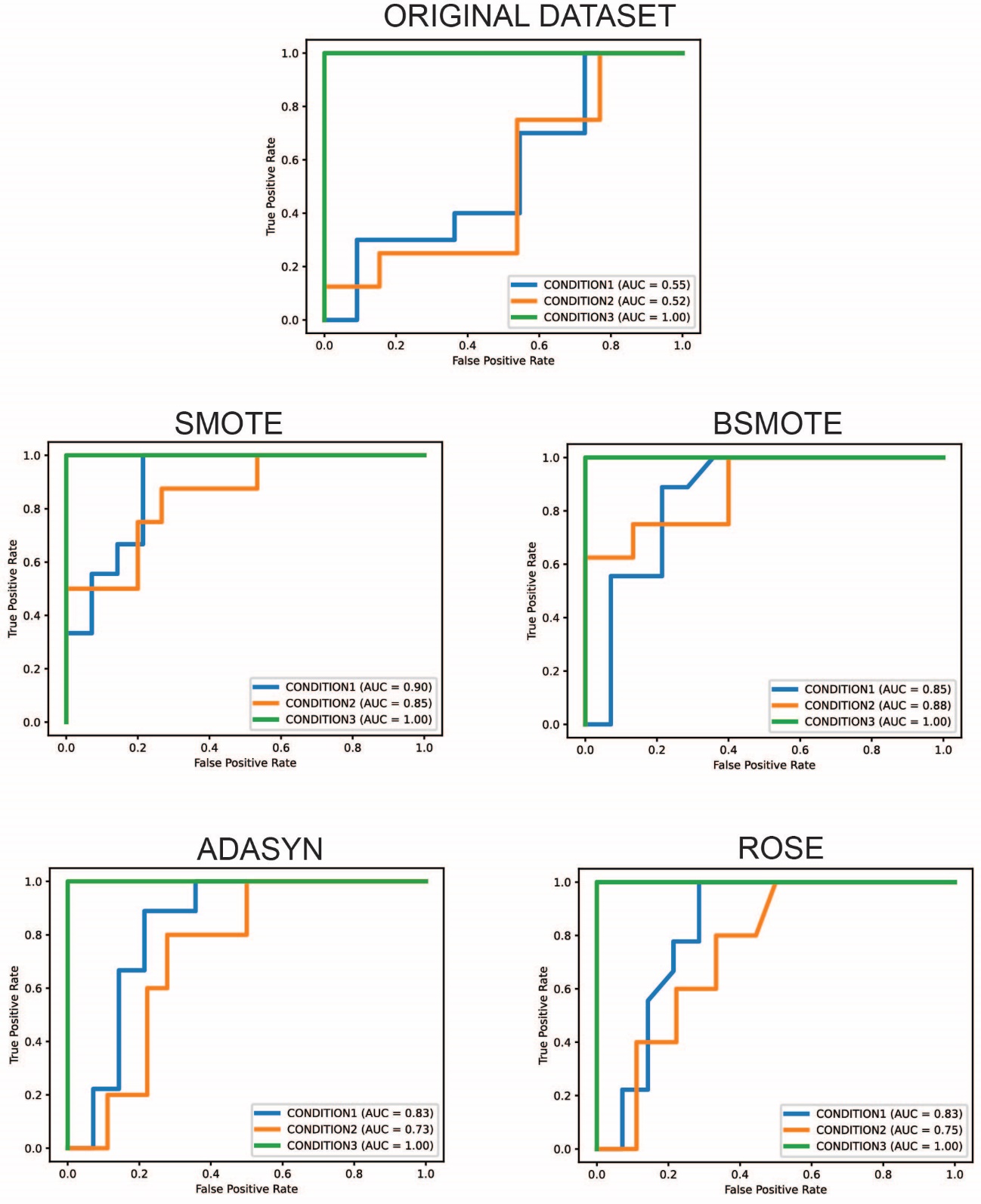

**Figure S2.** Comparison of PCA results using four different over-sampling methods all with z-score normalization of log transformed data. Effect of different over-sampling methods on PCA in this dataset is minor but none of the methods provided by META-BOA led to errors and grouping of synthetic samples.
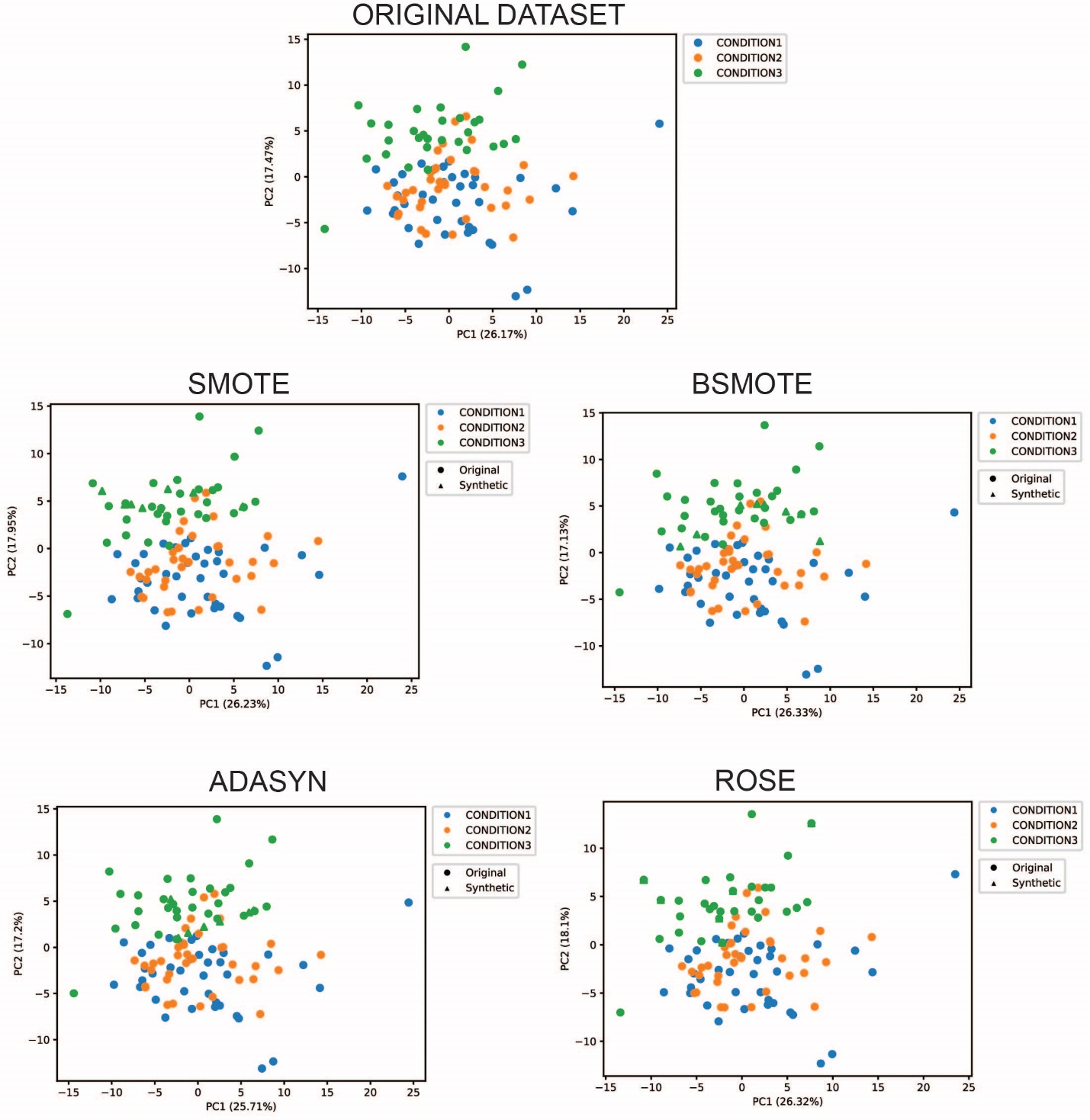

**Figure S3.** Comparison of t-SNE results using four different over-sampling methods all with z-score normalization of log transformed data. T-SNE once again shows that synthetic samples group with other samples from the same group and that four methods do not lead to major differences in projections.

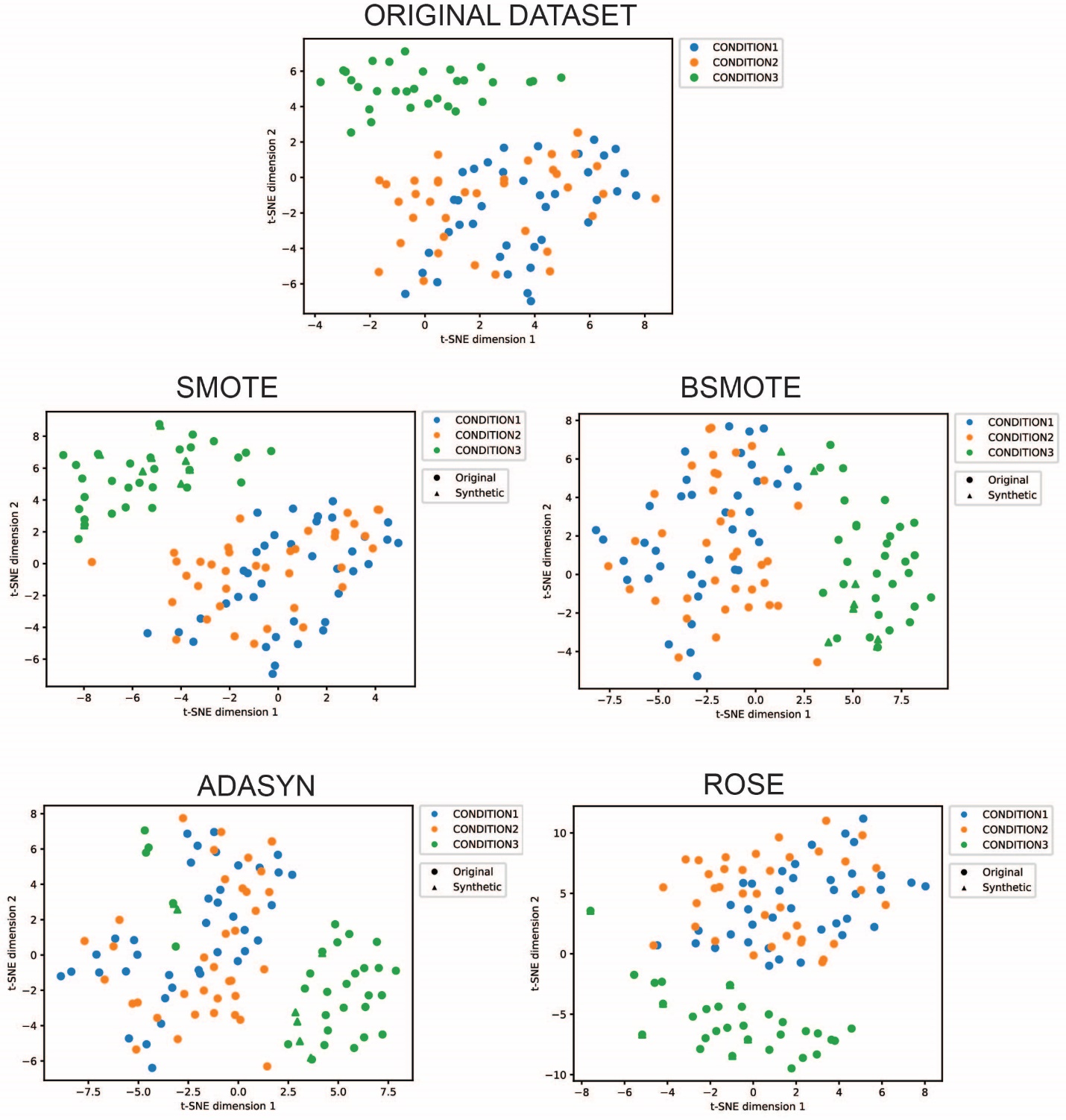
